# Enhanced sodium channel inactivation by temperature and FHF2 deficiency blocks heat nociception

**DOI:** 10.1101/2022.02.12.480202

**Authors:** Christopher Marra, Timothy V. Hartke, Matthias Ringkamp, Mitchell Goldfarb

## Abstract

Transient voltage-gated sodium currents are essential for the initiation and conduction of action potentials in neurons and cardiomyocytes. The amplitude and duration of sodium currents are tuned by intracellular fibroblast growth factor homologous factors (FHFs/iFGFs) that associate with the cytoplasmic tails of voltage-gated sodium channels (Na_v_s), and genetic ablation of *Fhf* genes disturbs neurological and cardiac functions. Among reported phenotypes, *Fhf2*^*null*^ mice undergo lethal hyperthermia-induced cardiac conduction block attributable to the combined effects of FHF2 deficiency and elevated temperature on the cardiac sodium channel (Na_v_1.5) inactivation rate. *Fhf2*^*null*^ mice also display a lack of heat nociception, while retaining other somatosensory capabilities. Here, we use electrophysiological and computational methods to show that the heat nociception deficit can be explained by the combined effects of elevated temperature and FHF2 deficiency on the fast inactivation gating of Na_v_1.7 and tetrodotoxin-resistant sodium channels expressed in dorsal root ganglion C fibers. Hence, neurological and cardiac heat-associated deficits in *Fhf2*^*null*^ mice derive from shared impacts of FHF deficiency and temperature towards Na_v_ inactivation gating kinetics in distinct tissues.

## 1. Introduction

Noxious cutaneous chemical, mechanical, and thermal stimuli open transient receptor potential (TRP) cationic channels at nociceptor afferent terminals,^30^ with resultant current influx and membrane depolarization triggering the activation of downstream voltage-gated sodium channels that mediate initiation and orthodromic conduction of action potentials.^16^ Loss-of-function mutations in genes encoding tetrodotoxin (TTX)-sensitive Na_v_1.7 and TTX-resistant Na_v_1.8 are hypoalgesic, reducing or ablating peripheral nociceptor activation by noxious stimuli,^1,3,7,13,21^ whereas gain-of-function missense mutations in these channel genes are hyperalgesic.^8,11,36^

The differential voltage-dependent gating properties of TTX-sensitive and TTX-resistant sodium channels collaborate to promote action potential initiation. Tetrodotoxin-sensitive channels, primarily consisting of Na_**v**_1.7,^6^ open more rapidly and at more negative membrane potential than does Na_v_1.8, but TTX-sensitive channels also inactivate at more negative voltages than Na_v_1.8.^2,6,9^ During a slow rise in voltage driven by stimulus-induced TRP channel currents at afferent terminals, most TTX-sensitive channels inactivate from the closed state, but residual channel availability provides a “primer” current depolarizing membrane sufficiently to open Na_v_1.8 channels that further drive action potential initiation.^5^ The orthodromic action potential rapidly depolarizes the downstream membrane, allowing for a greater contribution of Na_v_1.7 to spike conduction. Reduced availability of TTX-sensitive sodium channels near afferent terminals may explain why distal afferents harbor greater Na_v_1.8 density.^18^

Sodium channel gating is modulated by accessory proteins that include fibroblast growth factor homologous factors (FHFs, also termed intracellular fibroblast growth factors) bound to the carboxyl-terminal domain of Na_v_s.^14,19^ Most FHF isoforms induce a depolarizing shift in voltage dependence of Na_v_ fast inactivation without altering the voltage for activation,^10,20,33^ thereby increasing sodium current generated by Na_v_s. Mutations in FHF genes result in various disorders of excitable tissues.^12,15,22,23,25,28,29,31,34,35^ *Fhf2*^*null*^ mice are sensitive to ventricular cardiac conduction block when exposed to various stressors, including sodium or calcium channel blockers, uncoupling agents, or hyperthermia.^22,23^ FHF2 deficiency and elevated core body temperature collaborate to reduce availability and speed inactivation of cardiac Na_v_1.5 leading to failure of action potential conduction through myocardium.^22^

Genetic ablation of mouse FHF2 in peripheral neurons also results in the loss of cutaneous heat nociception.^35^ In this paper, we report that FHF2 modulates Na_v_1.7 and TTX-resistant sodium channel inactivation gating in a manner analogous to its effects on cardiac Na_v_1.5. Empirical and computational studies demonstrate that FHF2 modulation of Na_v_ fast inactivation provides a sodium current reserve in heated afferent terminals to enable action potential initiation and conduction. Overall, these data show that the limiting of sodium currents by FHF2 deficiency and elevated temperature constitutes a common gating mechanism underlying heat nociception and cardiac conduction deficits in *Fhf2*^*null*^ mice.

## 2. Materials and methods

### 2.1. Mouse genetics and genotyping

All work conducted with mice complied with protocol approved by the Hunter College Institutional Animal Care and Use Committee (IACUC). The murine embryonic stem cell clone EPD0339-4-F09 (International Mouse Phenotype Consortium) carrying a knockout first with conditional potential cassette bears loxP sites flanking (floxing) *Fhf2* exon 3 and Frt sites flanking the embedded G418-resistance and beta-galactosidase markers; these cells were used to derive mice carrying a *Fhf2*^*targeted*^ allele by blastocyst injection and outbreeding.^22^ The *Fhf2*^*targeted*^ allele was converted to an *Fhf2*^*nul1*^ allele by the injection of Cre recombinase-encoding plasmid into fertilized *Fhf2*^*targeted/+*^ eggs followed by reimplantation.^22^ Alternatively, the *Fhf2*^*targeted*^ allele was converted to the conditional *Fhf2*^*flox*^ allele by mating with mice that globally express a nuclear-targeted FLP recombinase ROSA26Sor^tm2(FLP*)Sor^ (Jackson Labs Stock #012930)^26^ to excise the embedded G418-resistance and beta-galactosidase markers. The *Fhf2*^*nul1*^ and *Fhf2*^*flox*^ alleles were backcrossed for 4 generations onto the 129S2 strain background. By crossing heterozygous *Fhf2*^*flox/+*^ females with male transgenic Advillin-Cre driver mice (Jackson Labs Stock #032536), which express Cre recombinase exclusively in the peripheral sensory neurons,^37,38^ *Fhf2*^*flox/Y*^*:Adv-Cre* test mice and *Fhf2*^*+/Y*^:Adv-Cre negative control littermates were generated. *Fhf2*^*flox*^ mice are publicly accessible through Jackson Labs (Stock #036280).

PCR genotyping was performed using PhireII Hot-Start DNA polymerase (Fisher Scientific, Waltham, MA). The *Fhf2*^*flox*^ and *Fhf2*^+^ alleles were distinguishable on PCR with primers IVS2-For (5’-GCCAGGAGTCTGCTCAACTCT) and IVS3-Rev (5’-GACTTTGGTGGGAGCATCCTGA), yielding products of 533 base pairs vs 388 base pairs, respectively.

### 2.2. Immunoblot detection of FHF2

Detergent-solubilized brain and dorsal root ganglion (DRG) tissue lysates were electrophoresed through 4% to 20% precast polyacrylamide gradient gel (Invitrogen, Waltham, MA), transferred to activated PVDF (Immobilon-P, Millipore Sigma, Burlington, MA), probed with rabbit anti-FHF2 polyclonal antibodies,^27^ and detected with peroxidase-conjugated secondary antibodies and enhanced chemiluminescence.

### 2.3. Sensory assays

All mice were conditioned to test apparati, environments, and modes of restraint for at least 2 consecutive days before data collection. *Tail-flick:* Heat nociception was evaluated using tail flick apparatus model 7360 (Ugo Basile, Trappe, PA). Mice were held with tail resting unrestrained on disk heat source, which on test initiation emitted infrared radiant heat 2 cm from the tip of the tail. The instrument measures the latency to tail withdrawal from the heat source. *Paw pinch:* Temperature-independent nociception was assayed using a force-calibrated forceps pressure transducer model Almemo 2450 (Ahlbom Mess, Holzkirchen, Germany) to determine exactly how much pressure applied to the hind paw of a mouse elicited a pain induced withdrawal response. Mice were placed in a tubular restraint with their hind limbs exposed. Force was applied to each hind paw, and the threshold response was recorded. *Von Frey filament assay:* Basal somatosensory response was measured using a set of sensory evaluator filaments (North Coast Medical, Morgan Hill, CA). Mice were placed in an elevated mesh-floored cage and allowed to explore and adapt for 20 to 30 minutes. Stimulation was manually applied to the hind footpads of the mouse starting with the thinnest softest filament and proceeding through filaments of greater thickness, performing 5 trials for each filament. Paw withdrawal was scored as a positive response, whereas bending of the filament without paw withdrawal constituted a negative response.

### 2.4. Voltage clamp analysis of Na_v_1.7 sodium currents in transfected HEK293 cells

Human embryonic kidney HEK293 cells (QBiogene) were transiently cotransfected with a 2:1 mixture of Na_v_1.7-expressing plasmid^8^ and a pIRES2-ZsGreen bicistronic plasmid (Takara Bio, San Jose, CA) expressing ZsGreen and mouse FHF2B proteins.^10^ The same pIRES2-ZsGreen plasmid without FHF2 coding sequence served as the control. Cells were trypsinized 3 hours posttransfection, seeded onto gelatinized coverslips, and were used for recording after 48 hours. For sodium current recordings, coverslips were transferred to recording the chamber containing carbogen-buffered bath solution (115 mM NaCl, 26 mM NaHCO_**3**_, 3 mM KCl, 10 mM glucose, 4 mM MgCl_2_, 2 mM CaCl_2_, 0.2 mM CdCl_2_, 3 mM myoinositol, 2 mM Na pyruvate, 7 mM NaOH-buffered HEPES, and pH 7.2) at 25°C and green fluorescent cells were whole-cell patched with pipettes filled with 104 mM CsF, 50 mM tetraethylamine chloride, 10 mM HEPES pH 7.2, 5 mM glucose, 2 mM MgCl_2_, 10 mM EGTA, 2 mM ATP, and 0.2 mM GTP and having 1 to 2 MΩ resistance. All voltage commands and current recordings were made using a MultiClamp 700 amplifier, digital or analog converter Digidata 1440, and pClamp10 software (Molecular Devices, San Jose, CA). For all recording protocols, voltage-gated current was isolated during data acquisition by P/N subtraction of leak and capacitive currents (N = −6).

#### 2.4.1. Channel activation protocol

For activation measurements, voltage was stepped from −110 mV to reporting voltages ranging from −60 mV to 0 mV in 5 mV increments. As the criterion for the adequate clamp, transient current peaks for all voltage commands were nested within the larger current trace of a following or preceding voltage step command.

#### 2.4.2. Steady-state channel inactivation protocol

For inactivation measurements, 80 milliseconds conditioning voltages ranging from −110 mV to −50 mV in 5 mV increments were followed by 10 milliseconds 0 mV reporting voltage to open noninactivated channels.

#### 2.4.3. Voltage ramp protocol

As a measure of a closed-state channel inactivation rate, a 10-sweep protocol used depolarization from −110 mV to −20 mV either instantaneously (step) or as a ramp ranging from 2 to 18 milliseconds in 2 milliseconds intervals. After recording at 25°C, temperature was elevated to 35°C and later to 40°C. At elevated temperatures, the cell was first tested as abovementioned to ensure the maintenance of tight clamp, after which the voltage ramp protocol was conducted. For many cells, voltage ramp protocols could be successfully run at all 3 temperatures.

### 2.5. Voltage clamp analysis of tetrodotoxin-resistant sodium currents in acutely dissociated DRG neurons

DRGs were plucked from spinal columns of 8-to-12-week-old mice and trimmed free of associated nerves. Neurons were dissociated by treatment with 1% trypsin and 150 μg/mL collagenase in Hank balanced salt solution at 37°C for 30 minutes followed by washing into Dulbecco modified Eagle medium (DMEM) + 10% fetal bovine serum (FBS) +250 μg/mL DNasel and trituration with fire-polished Pasteur pipettes. Dissociated cells were washed 3 times with DMEM + 10% FBS and seeded on to 12 mm diameter coverslips precoated with poly-D-lysine and laminin at a density of 3 × 10^4^ cells/coverslip and cultured for 18 to 40 hours before recording. Recordings were conducted in the extracellular solution consisting of 24 mM NaCl, 3 mM KCl, 3 mM CsCl, 26 mM NaHCO_3_, 50 mM choline Cl, 11 mM glucose, 25 mM TEA-Cl, 2 mM Na pyruvate, 3 mM myoinositol, 10 mM HEPES, 5 mM 4-aminopyridine, 2mM MgCl_2_, 1 mM BaCl_2_, 1 mM CaCl_2_, and 0.2 mM CdCl_2_ and 0.5 μM TTX at pH 7.2 using patch pipettes of 1 to 1.5 MΩ resistance when filled with 120 mM K gluconate, 4 mM NaCl, 5-mM KOH-buffered HEPES, 5-mM KOH-buffered EDTA, 15 mM glucose, 1 mM MgSO_4_, 3 mM Mg-ATP, and 0.1 mM Na_2_-GTP at pH 7.2. The voltage dependence of TTX-resistant sodium channel steady state inactivation was assayed in a 13-sweep protocol using 80 milliseconds conditioning voltage ranging from −60 mV to 0 mV in 5 mV increments followed by + 10 mV reporting voltage.

### 2.6. Temperature-induced excitation of acutely dissociated DRG nociceptive neurons

#### 2.6.1. Voltage-sensitive dye recordings

DRG neurons grown on coverslips as described above were first treated with a blocking solution consisting of sterile HBSS with 10 mM HEPES + 1% horse serum for 5 minutes at room temperature followed by staining with HBSS/HEPES + 70 mM Di-8-ANEPPS (Biotium, Fremont, CA) for 15 minutes room temperature and recovery in HBSS/HEPES + 10% horse serum for 5 minutes. Coverslips of stained neurons were transferred to the recording chamber containing carbogen-bubbled extracellular solution: 115 mM NaCl, 26 mM NaHCO_3_, 3 mM KCl, 1.2 mM KH_2_PO_4_, 3 mM glucose, 2 mM myoinositol, 2 mM Na pyruvate, 7 mM HEPES, 1.2 mM MgSO_4_, 2 mM CaCl_2_, and 0.2 mM CdCl_2_ at pH 7.2. Using a 40X lens objective, SOLA-SE LED illumination (Lumencor, Beavorton, OR) and a filter cube consisting of a 520 ± 45 nm excitation interference filter, a dichroic mirror with 570 nm central wavelength, and 610 nm barrier emission filter, fluorescence images were captured at 2 kHz with a NeuroCCD-SM camera (Redshirt Imaging, Decatur, GA) while heating the bath from 32°C to 44°C at a rate of approximately 4°C per minute. Action potentials were detected as transient fluorescence changes of 2% to 3% resting fluorescence. Excitable cells were then imaged by high-resolution bright field illumination to assess the neuronal cell diameter.

#### 2.6.2. Current-clamp recordings

Recordings were performed using the same extracellular solution as abovementioned and patch pipettes of 2 to 3 MΩ resistance when filled with 120 mM K gluconate, 4 mM NaCl, 5-mM KOH-buffered HEPES, 5-mM KOH-buffered EDTA, 15 mM glucose, 1 mM MgSO_4_, 3 mM Mg-ATP, and 0.1 mM Na_2_-GTP at pH 7.2. Cells approximately 12 μm in diameter were selected for patching based on prior optical imaging that had determined the size of heat responsive cells. Excitability was first assessed at 32°C by applying 200 millisecond current pulses ranging from 0 to 400 pA in 25 pA intervals. Excitable cells were then held with 0 pA current injection while heating the chamber to 45°C at approximately 4°C per minute. Voltages were recorded at 10 kHz acquisition rate.

### 2.7. Temperature-induced modulation of C-fiber compound action potential conduction

Saphenous nerve segments approximately 10 mm in length were used for recording C-fiber compound action potentials (CCAPs) as previously described.^18^ Each end of a nerve was drawn into a glass pipette and electrically isolated from the central nerve with petroleum jelly to establish high-resistance electrical seals. The central nerve was bathed in continuously replenished carbogen-bubble synthetic interstitial fluid. One nerve end was stimulated with a 100 microsecond current pulse every 9 seconds, whereas a recording electrode placed at the other end of the nerve monitored evoked potentials. Recordings were initiated at 27°C with the stimulus strength adjusted sufficient to yield a maximal evoked response. Thereafter, a heating element in the buffer flow path was used to heat the bath up to 45°C at a rate of approximately 0.2°C per second and then cooled back to 27°C at the same rate. Only CCAPs from the declining temperature ramp were used for analysis.

### 2.8. Computational modeling of wild-type and Fhf2^null^ nociceptor excitation

A computational model of a nociceptive neuron with realistic afferent terminal branching and TTX-sensitive and—resistant transient voltage-gated sodium conductances constructed on the NEURON simulation platform^4,17^—was adapted for our construction of temperature-responsive wild-type and *Fhf2*^*nul1*^ nociceptor models. For all gated ion conductance models within the neuron, we applied a rate scaling factor Q10 of N^^^log_10_[localtemp(°C)—37], where N ranged from 2.3 to 3.0 and where localtemp is the temperature in a particular segment of the neuron, thereby allowing for localized temperature modulation. The *Fhf2*^*null*^ nociceptor model was identical to the wild-type model, except for altering the parameters within the TTX-sensitive and—resistant sodium conductances to induce left shifts in V_1/2_ inactivation and accelerated inactivation to an extent consistent with experimental findings. Another modification in the models was to increase TTX-resistant density in the branched terminal afferents, consistent with empirical data.^18^ These wild-type and *Fhf2*^*null*^ nociceptor models will be deposited at the publicly accessible repository ModelDB (https://senselab.med.yale.edu/modeldb/).

To assess axonal conduction of action potentials, a segment of the main axon was stimulated with a 100 microsecond 400 pA current pulse while setting the entire neuron to either 37°C or 43°C. Transient receptor potential channel activation was simulated as inward current ramps delivered into all termini of the afferent tree. For simulation of mechanosensation, the entire neuron was maintained at 37°C, whereas for simulation of heat nociception, all segments of the afferent tree within 250 μm of termini were raised to 43°C before injection of current ramps. To simulate heat stimulation of an acutely dissociated model neuron, the axon was disconnected from the soma, the density of TTX-resistant conductance was increased to that found at branched termini, and a current ramp was injected into the soma set to 45°C.

## 3. Results

### 3.1. FHF2 deficiency in sensory neurons causes loss of heat nociception with lesser effects on other somatosensory modalities

*Fhf2*^*null*^ mice *(Fhf2*^*−/Y*^ male and *Fhf2*^*−/−*^ female) maintained on the 129S2 background strain are viable and fertile but are sensitive to cardiac conduction block when core body temperature rises above 41°C.^22^ When *Fhf2*^*−/Y*^ and *Fhf2*^*−/−*^ mice were tested for heat nociception by the tail flick assay, they were uniformly nonresponsive over 12 seconds exposure of the tail to the infrared heating beam, whereas wild-type *Fhf2*^*+/Y*^ and *Fhf2*^*+/+*^ mice sensed heat and flicked their tails in 3.2 ± 0.3 seconds (**Fig. 1A**). To determine whether heat nociception requires FHF2 expression in nociceptive neurons, *Fhf2*^*flox/+*^ mice heterozygous for a loxP-flanked (*flox*) *Fhf2* conditional allele (see Methods) were mated with Advillin-Cre transgenic mice that express Cre recombinase exclusively in peripheral neurons.^37,38^ Male *Fhf2*^*flox/Y*^*:AdvCre* progeny showed complete absence of FHF2 protein in DRG neurons while retaining normal FHF2 levels in the brain (**Fig. 1B**). When tested in the tail flick assay, *Fhf2*^*flox/Y*^: *AdvCre* mice were uniformly nonresponsive, whereas control animals retaining peripheral FHF2 expression (*Fhf2*^*flox/Y*^ and *Fhf2*^*+/Y*^*:AdvCre*) responded to heat stimulus in 3.0 ± 0.2 seconds (**Fig. 1C**).

**Figure 1.**
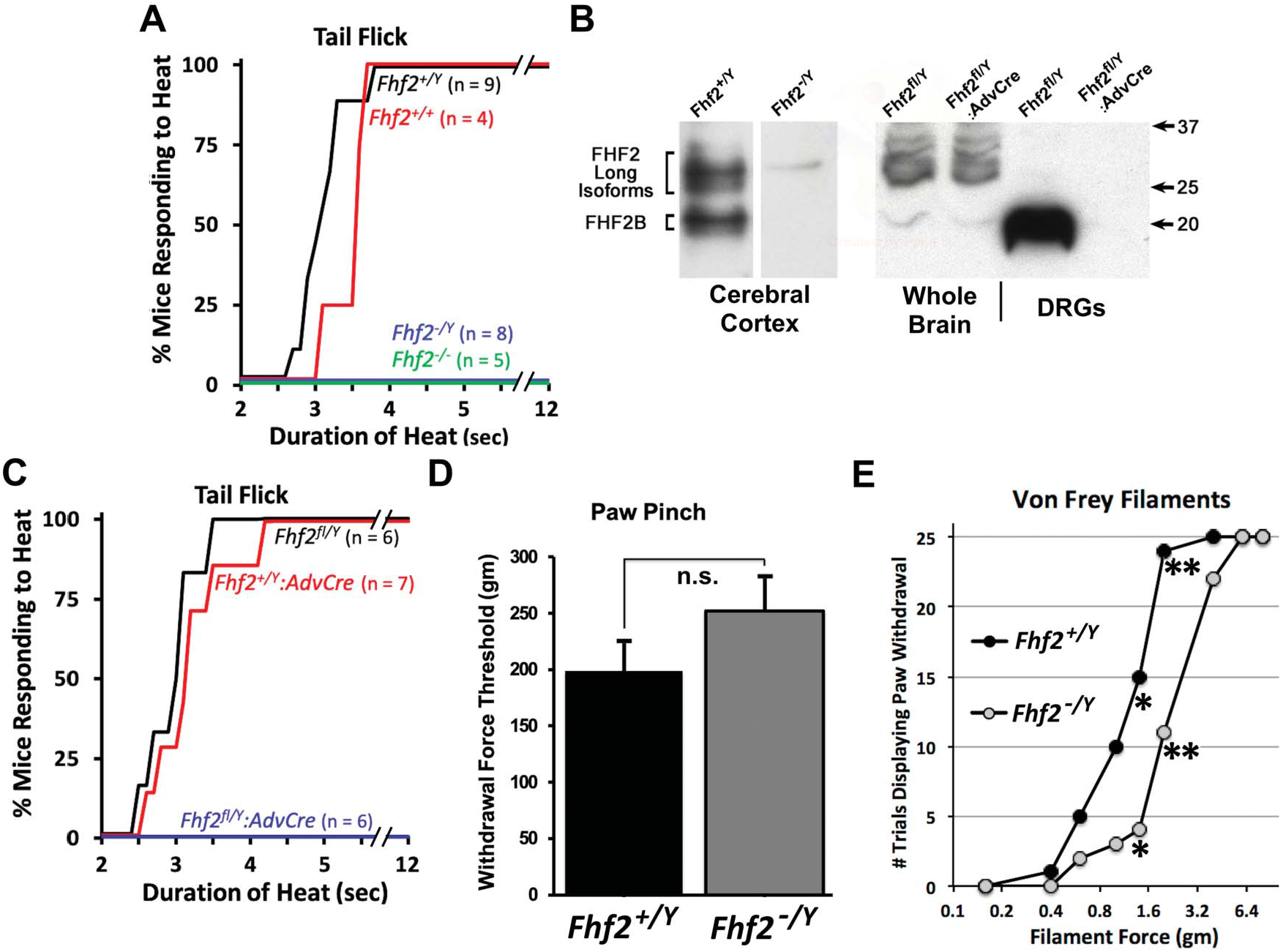
Somatosensation in mice harboring global or peripheral neuron-specific *Fhf2* gene ablation. (A) Tail flick assay of heat nociception in wild-type and *Fhf2*^*null*^ mice. Cumulative percentage plot for tail flick as function of tail heating time for wild-type female (*Fhf2*^*+/+*^), wild-type male (*Fhf2*^*+/Y*^*), Fhf2*^*null*^ female (*Fhf2*^*-/-*^), and *Fhf2*^*null*^ male (*Fhf2*^*-/Y*^) mice. Male and female *Fhf2*^*null*^ mice were not responsive to heating. n, number of mice. (B) FHF-2 protein expression in wild-type and mutant mice. Postnuclear protein extracts from whole brain, cerebral cortex, and DRGs were subjected to SDS polyacrylamide gel electrophoresis and immunoblotting with antibodies directed against the C terminus of FHF2 shared by all FHF2 isoforms. Lower molecular weight FHF2 (FHF2B) and higher molecular weight FHF2 isoforms were present in wild-type, but not *Fhf2*^*null*^, cerebral cortex (left). Male mice carrying a floxed *Fhf2* allele (*Fhf2*^*fl/Y*^) and the Advillin-Cre transgene (*Adv-Cre*) showed complete loss of FHF2B expression in DRG, while retaining FHF2 expression in the brain. (C) Tail flick assay in mice with DRG-specific ablation of FHF2 expression. Cumulative percentage tail flick plot shows that mice with DRG-specific ablation of FHF-2 expression (*Fhf2*^*fl/Y*^*:Adv-Cre*) were insensitive to heating, whereas control mice (*Fhf2*^*+/Y*^*:Adv-Cre* and *Fhf2*^*fl/Y*^) all rapidly responded to tail heating. n, number of mice. (D) Paw pinch assay of mechanical nociception in wild-type and *Fhf2*^*nul1*^ mice. For each animal, the footpad was squeezed with a force-calibrated forceps until a withdrawal response was elicited. n.s., not significant impairment of mechanical nociception. (E) Von Frey filament assay of touch sensation in wild-type and *Fhf2*^*null*^ mice. Five male wild-type (*Fhf2*^*+/Y*^) and mutant (*Fhf2*^*-/Y*^) mice were placed without restraint on a mesh platform, a hind paw footpad was touched with nylon filaments of graded thickness or force, and sensation was scored by paw withdrawal. Five trials were conducted for each filament per mouse (25 trials per genotype cohort). Graph plots number of trials (of 25) that a response was elicited. *\*P <* 0.004; *\*\*P <* 0.0003. FHF, factor homologous factor.

*Fhf2*^*nul1*^ and wild-type mice were tested for mechanical nociception by the paw pinch assay, which entails slowly increasing the compression force on the footpad until a response is elicited. *Fhf2*^*+/Y*^mice responded to compression at a force of 199 ± 25 gm (n = 7), whereas *Fhf2*^*-/Y*^ mice responded at 252 ± 30 gm (n = 9; *P <* 0.21) (**Fig. 1D**). Therefore, *Fhf2*^*nul1*^ mice retain relatively near-normal mechanosensitivity. *Fhf2*^*nul1*^ and wild-type mice were also tested for paw sensitivity to touch using graded diameter von Frey nylon filaments. *Fhf2*^*-/Y*^ mice were moderately but significantly less sensitive to touch compared with *Fhf2*^*+/Y*^ littermates (**Fig. 1E**). In summary, *Fhf2*^*null*^ mice have multiple somatosensory deficits, with heat nociception completely absent, and FHF2 is functionally required in heat sensitive, nociceptive neurons. Our findings are consistent with those reported previously for nociception when peripheral neurons were rendered *Fhf2*^*null*^ on a different background strain.^35^

### 3.2. FHF2 modulation of inactivation gating for sodium channels expressed in nociceptive neurons

Na_v_1.7 is the most abundantly expressed sodium channel in nociceptive neurons,^5,6^ and the Na_v_1.7 cytoplasmic tail has been shown to bind FHF2.^35^ Indeed, the crystal structure of FHF2 in complex with the tail of cardiac Na_v_1.5 revealed that the channel residues that interface with FHF2 are conserved within all 9 mammalian sodium channel alpha subunits.^32^ To assess potential effects of FHF2 on Na_v_1.7 inactivation gating, HEK293 cells (from QBiogene) were cotransfected with Na_v_1.7 expression plasmid^8^ together with either an FHF2B/GFP bicistronic expression plasmid or GFP-only plasmid.^10^ Fluorescent cells were selected for patching and voltage clamp recording of sodium currents (see Methods). The short B isoform of FHF2 protein was chosen for our analyses based on immunoblotting data showing that FHF2B predominates in peripheral neurons, whereas higher molecular weight FHF2 species are more abundant in the brain (**Fig. 1B**) and heart.^22^

The voltage dependence of steady-state Na_v_1.7 inactivation was V_1/2_ = −86.8 ± 1.0 mV in the absence of FHF2B (n = 11) and was right-shifted to V_1/2_ = −71.6 ± 1.8 mV when FHF2B was present (n = 11, *P <* 0.000002) (**Figs. 2A and B**). By contrast, FHF2B did not significantly alter the voltage dependence of Na_v_1.7 activation (**Figs. 2A and B**). Furthermore, FHF2B did not significantly alter Na_v_1.7 peak current density measured at −20 mV: w/o FHF −77 ± 19 pA/pF (n = 11), with FHF2B −88 ± 13pA/pF (n 5 11), *P* > 0.6.

**Figure 2.**
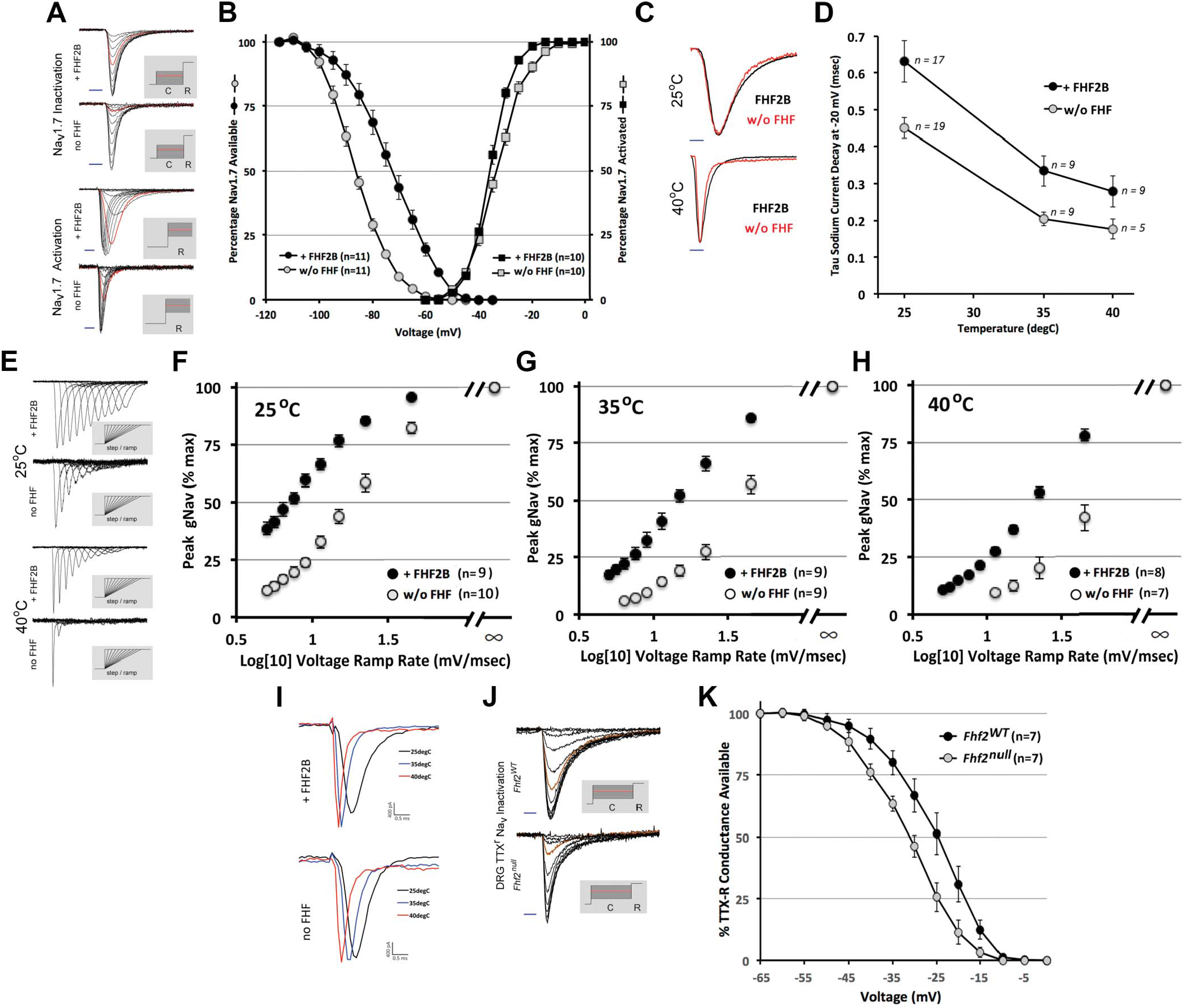
FHF-2 modulation of sensory neuronal sodium channel inactivation. (A and B) Voltage dependence of Na_v_1.7 steady-state inactivation and activation with or without FHF2B at 25°C in HEK cells. (A) Exemplar superimposed Na_v_1.7 sodium current traces ±FHF2B induced by voltage-clamp protocols (shaded inserts) for steady-state inactivation (upper) and activation (lower). For inactivation measurements, 80 milliseconds conditioning voltages, C, ranging from −110 mV to −50 mV (−70 mV shown in red) were followed by 10 milliseconds 0 mV reporting voltage, R, to open noninactivated channels. For activation measurements, voltage was stepped from −110 mV to reporting voltages, R, ranging from −60 mV to 0 mV (−30 mV shown in red). Blue scale bars, 1 milliseconds. (B) The presence of FHF2B induced a 15 mV depolarizing shift in V_1/2_ inactivation (*P <* 0.000002) without significantly impacting activation. n, number cells analyzed. (C) Normalized Na_v_1.7 current traces with or without FHF2Bat 25°C and 40°C. (D) Na_v_1.7 current decay time constants at −20 mV ±FHF2Bat 25°C, 35°C, and 40°C. FHF2B modestly but significantly slows channel inactivation at all temperatures (P < 0.05) (E-H) Na_v_1.7 peak conductance on ramped vs step depolarization ±FHF2 at different temperatures. (E). Exemplar superimposed Na_v_1.7 sodium current traces ±FHF2Bat 25°Cand 40°C induced by depolarization from −110 mV to −20 mV instantaneously (step) or as a ramp ranging from 2 to 18 milliseconds in 2 milliseconds intervals (shaded inserts). Graphed representation of all cell data at (F) 25°C, (G) 35°C, and (H) 40°C. Slower voltage ramp rates reduce sodium channel opening via by-state inactivation, with much greater reduction in the absence of FHF2B (P < 0.0003 with vs without FHF for all ramp rates and temperatures). (I) Na_v_1.7 current amplitudes at −20 mV at different temperatures. An HEK cell expressing Na_v_1.7 and FHF2B (upper traces) or Na_v_1.7 only (lower traces) was recorded at 25°C, 35°C, and 40°C. In neither cell did the peak sodium currents significantly diminish at elevated temperature. (J, K) Steady-state inactivation of TTX-resistant sodium channels in wild-type and *Fhf2*^*null*^ DRG neurons. (J) Exemplar superimposed TTX-resistant sodium current traces in wild-type (upper) and *Fhf2*^*null*^ (lower) DRG neuron induced using steady-state inactivation protocol (shaded inserts) consisting of 80 milliseconds conditioning voltage, (C), ranging from −60 mV to 0 mV in 5 mV increments (−25 mV shown in red) followed by 10 mV reporting voltage. Blue scale bars, 2.5 milliseconds. (K) V_1/2_ inactivation of DRG TTX-resistant sodium current undergoes a 6 mV hyperpolarizing shift in *Fhf2*^*null*^ neurons compared with *Fhf2*^*WT*^ (*P <* 0.02). FHF, factor homologous factor; Na_v_, voltage-gated sodium channels; TTX, tetrodotoxin; WT, wild-type.

We then examined the Na_v_1.7 fast inactivation rate from the open state at 25°C, 35°C, and 40°C in the presence vs absence of FHF2B on depolarization to −20 mV. At 25°C, Na_v_1.7 inactivated somewhat more slowly in the presence of FHF2B (n = 17, τ = 0.63 ± 0.05 ms) vs its absence (n = 19, τ = 0.45 ± 0.03 ms, *P <* 0.01, **Figs. 2C and D**). Na_v_1.7 inactivated progressively faster at 35°C and 40°C, but at each temperature, inactivation was significantly slowed by the presence of FHF2B (**Figs. 2C and D**). Hence, the shortest duration of Na_v_1.7 current was measured at 40°C in the absence of FHF2B (n = 5, τ = 0.18 ± 0.03 ms, **Figs. 2C and D**) because of the independent effects of elevated temperature and FHF vacancy.

The rate of closed-state Na_v_1.7 inactivation was assayed at different temperatures in the presence vs absence of FHF2B by a voltage ramp protocol, as previously described.^21^ Voltage was raised from −110 mV to −20 mV either instantaneously (step) or linearly over times ranging from 2 to 18 milliseconds (ramps), and the Na_v_1.7 peak sodium conductance for each voltage ramp was expressed as percentage of the peak conductance that had been achieved by the direct voltage step. Under each transfection and temperature condition, Na_v_1.7 peak conductance diminished as voltage ramp speed was reduced (**Figs. 2E–H**), reflecting increased closed-state inactivation. At any voltage ramp speed, inactivation was dramatically enhanced by both elevated temperature and the absence of FHF2B (*P* < 0.0003) (**Figs. 2E–H**). For example, 68 ± 5% of channels underwent closed-state inactivation during a rapid 45 mV/ms ramp at 40°C in the absence of FHF2B (**Figs. 2E and H**), whereas this ramp speed and temperature induced inactivation of only 22 ± 3% of channels in the presence of FHF2B (**Figs. 2E and H**, *P <* 0.0003).

It has been reported that Na_v_1.7 undergoes internalization at elevated temperatures in the absence of FHF2 and that reduced Na_v_1.7 density underlies the heat nociception deficit of Fhf2-deficient mice.^35^ In our recordings, we failed to see a consistent reduction of Na_v_1.7 peak current at elevated temperature in the absence or presence of FHF2B (**Fig. 2I**). Therefore, the major mechanism for heat-induced Na_v_1.7 current reduction appears to be accelerated channel inactivation in the absence of FHF.

In small-diameter nociceptive neurons, Na_**v**_1.7 is the principal TTX-sensitive sodium channel, whereas Na_v_1.8 and, to a lesser extent, Na_v_1.9 are responsible for the transient TTX-resistant sodium current. Transient TTX-resistant sodium currents in acutely dissociated wild-type and *Fhf2*^*null*^ DRG neurons were analyzed by voltage clamp. Steady-state inactivation occurred with V_1/2_ = −31 mV (n = 7) in wild-type neurons vs V_1/2_ = −24.5 mV (n = 7, *P <* 0.025) in *Fhf2*^*null*^ neurons (**Figs. 2J and K**), demonstrating that FHF2 modulates inactivation gating of all sodium channel isoforms in nociceptive neurons.

### 3.3. FHF2 is not required for heat-induced excitation of acutely dissociated DRG neurons

FHF2 is required in peripheral neurons for heat nociception and FHF2 deficiency facilitates inactivation of the Na_v_s expressed in DRGs. Therefore, we first sought to analyze heat-induced excitation of acutely dissociated *Fhf2*^*WT*^ and *Fhf2*^*null*^ DRG neurons.

To determine which cell sizes in the heterogeneous DRG population to focus on for patch clamp analysis, we first surveyed fields of adult acutely dissociated *Fhf2*^*WT*^ neurons for heat-induced fluorescence spikes following extracellular staining with voltage-sensitive dye Di8-ANEPPS. In one visual field containing several stained cells (**Fig. 3A**), one of 4 cells commenced firing action potentials on heating to 43°C (**Fig. 3B**). This cell (**Fig. 3C**) and 2 other optically identified heat excitable neurons among 37 cells screened in other fields had diameters of approximately 12 μm.

**Figure 3.**
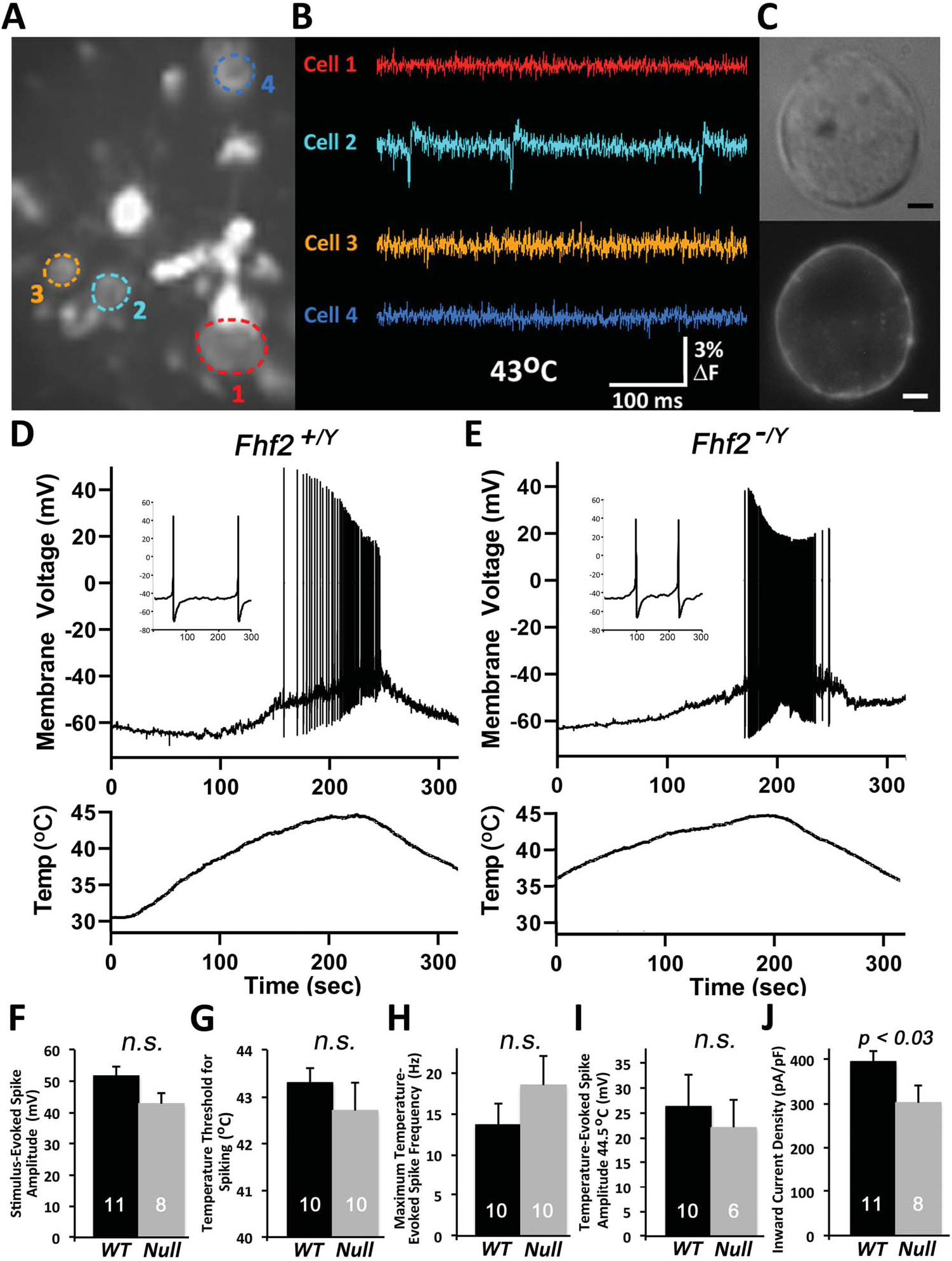
Excitation by heat of acutely dissociated DRG neurons. (A–C) Optical detection of heat-sensitive nociceptor excitation. Acutely dissociated wild-type DRG neurons were cultured overnight and stained with di-8-ANEPPS. A visual field of 4 cells was imaged with a high speed CCD camera while heating to 43°C. (A) A single image shows fluorescence of neurons (color circled) along with debris including myelin. (B) Fluorescence intensity monitored at 2 kHz shows action potentials generated in cell 2. (C) Bright field and fluorescent high magnification imaging of cell 2. Scale bar, 2 mm. (D) Voltage recording of a patched 12 mm diameter *Fhf2*^*WT*^ neuron during heating and cooling. Inset is an expanded time scale showing 2 action potentials early in the spike train. (E) Voltage recording of a patched and heated *Fhf2*^*null*^ neuron. (F-J) Excitation and current properties of *Fhf2*^*WT*^vs *Fhf2*^*null*^ heat sensitive cells. (F) Current stimulus-evoked spike amplitude at 32°C, (G) temperature threshold for spike initiation, (H) maximum temperature-evoked spike frequency, (I) spike amplitude at 44.5°C, and (J) inward current density on voltage-clamp step from −70 mV to 0 mV at 32°C. Number of recorded cells and standard errors are indicated for each recording parameter. FHF, factor homologous factor; n.s., not significant; WT, wild-type.

For electrical recordings, *Fhf2*^*WT*^ DRG neurons were dissociated, cultured overnight, and ∼12 μm diameter cells selected for patching. Patched cells were first analyzed in the voltage clamp for inward sodium and outward potassium currents in response to command depolarization, followed by excitability in response to inward current injection. Current-excitable cells were then subjected to temperature ramping from 32°C to 45°C followed by cooling while recording voltage. **Figure 3D** shows an example of a heat-excitable *Fhf2*^*WT*^ neuron which commenced firing at 42°C. **Figure 3E** shows heat-induced excitation of an *Fhf2*^*null*^ neuron. The fraction of *Fhf2*^*WT*^ neurons that were excitable by heating (39 of 96 cells) was statistically indistinguishable from the fraction of heat-excitable *Fhf2*^*null*^ neurons (13 of 31 cells). By a range of criteria, including current stimulus-evoked spike amplitude (**Fig. 3F**), temperature threshold for spike induction (**Fig. 3G**), maximum temperature-evoked spike frequency (**Fig. 3H**), and temperature-evoked spike amplitude (**Fig. 3I**), *Fhf2*^*null*^ neurons were as excitable as their wild-type counterparts. The only parameter significantly affected by *Fhf2* deletion was a small reduction in the amplitude of peak inward current on voltage-clamp depolarization from −70 mV to 0 mV (**Fig. 3J**). This likely reflects a hyperpolarizing shift in V_1/2_ steady state inactivation of Na_v_1.7, rendering some of the channels inactivated at −70 mV before depolarization.

We suspect that acutely dissociated *Fhf2*^*nul1*^ DRG neurons retain heat-induced excitability due to the transient reorganization and expression of nociceptor-gated and voltage-gated channels after the severing of axonal processes (see Discussion).

### 3.4. Rapid and reversible heat-induced conduction failure in Fhf2^null^ C-fibers

If altered voltage dependence and kinetics of sodium channel inactivation in *Fhf2*^*nul1*^ nociceptors account for loss of heat nociception, it should be possible to detect a rapidly reversible impairment in *Fhf2*^*nul1*^ sensory action potential generation or conduction on heating. Towards this end, we tested the ability of unmyelinated sensory C-type fibers to conduct action potentials at different temperatures. Electrical stimulation of *Fhf2*^*WT*^ and *Fhf2*^*nul1*^ saphenous nerves ex vivo^18^ generated CCAPs (**Figs. 4A and B**). Each nerve was then stimulated every 9 seconds while being warmed to 45°C and cooled back to 27°C, and CCAPs from the cooling phase of the temperature ramp were analyzed. Elevated temperature increased CCAP conduction velocity and decreased amplitude (**Figs. 4C–F**), reflecting faster rates of voltage-gated channel activation and inactivation. In *Fhf2*^*WT*^ nerves, CCAP amplitudes declined linearly from 27°C to 44°C (**Figs. 4C,E,F**). C-fiber compound action potential amplitudes in *Fhf2*^*null*^ nerves showed a similar decrement up to 40°C, but declined precipitously above 40°C (**Figs. 4D–F**), reflecting conduction block or severe attenuation in at least half of all mutant fibers. **Figures 4D and E** panels illustrate that *Fhf2*^*null*^ CCAP impairment was reversed within 27 seconds (3 stimuli) as nerves were cooled from 44°C to 40°C.

**Figure 4.**
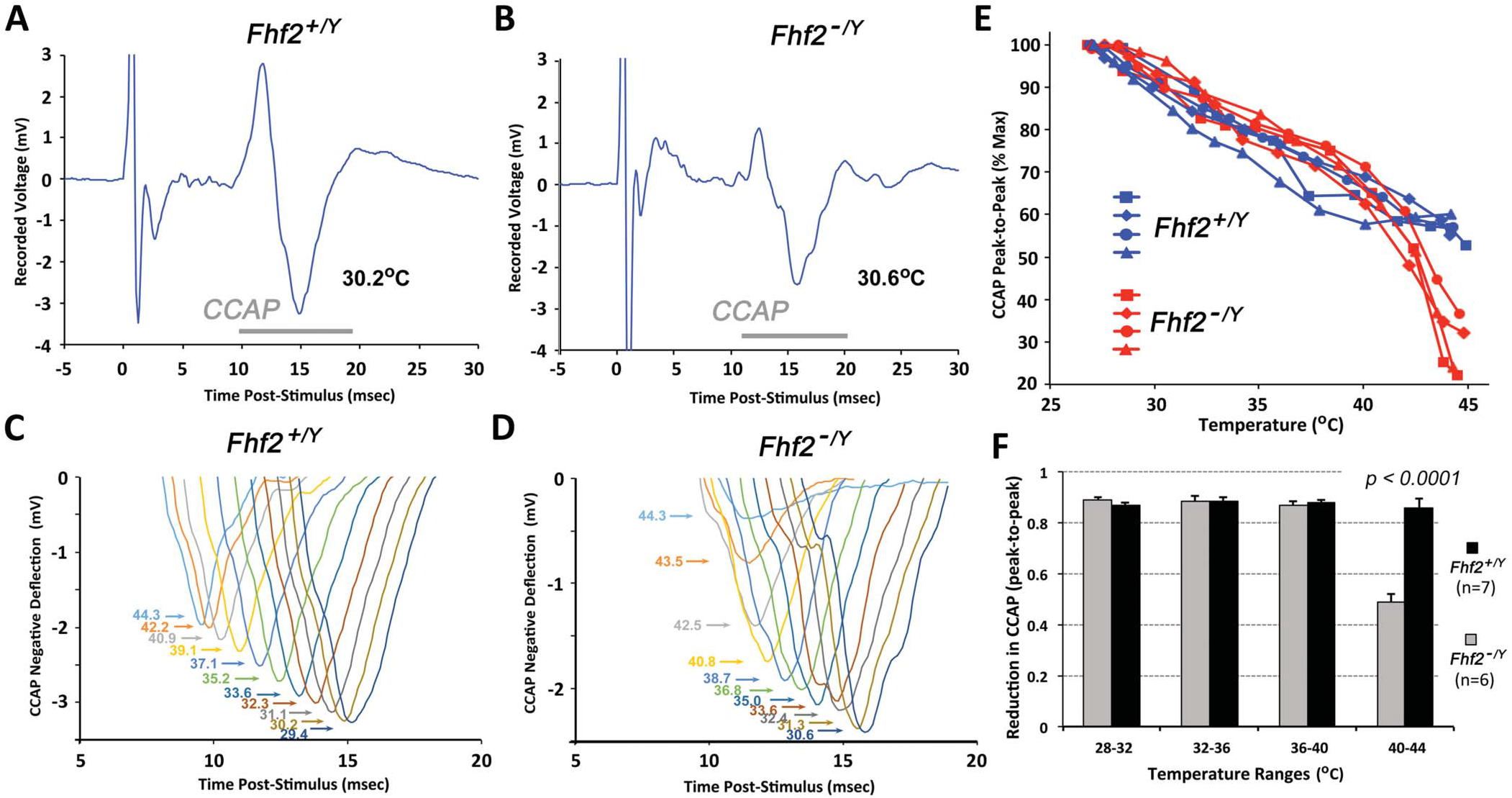
Temperature modulation of C-fiber compound action potential conduction. (A) Stimulus response of the wild-type saphenous nerve. The nerve maintained at 30 to 31°C was stimulated at one end at time - 0 milliseconds. The recording electrode detected an instantaneous stimulus artifact, fast arriving voltage deflections (1-3 ms) reflecting activation and conduction of myelinated axons, and slow C-fiber compound action potential (CCAP) from 10 to 20 milliseconds poststimulus. (B) Stimulus response of *Fhif2*^*null*^ saphenous nerve. A similar CCAP was detected at 30 to 31°C. (C) Superimposed CCAP traces of a wild-type nerve stimulated every 9 seconds during cooling from 44.3°C down to ∼30°C. CCAPs displayed a near-linear increase in amplitude and decrease in velocity across the temperature range. (D) Superimposed CCAP traces of an *Fhf2*^*null*^ nerve stimulated every 9 seconds during cooling from 44.3°Cdown to ∼30°C. The amplitude ofthe CCAP negative deflection fell precipitously when nerve temperature exceeded 41°C. (E) Plot of CCAP amplitudes vs temperature. For each of 4 wild-type and 4 *Fhf2*^*null*^ nerves, the positive-to-negative peak-to-peak CCAP amplitudes at each temperature were plotted as a percentage of the amplitude recorded at 27°C. Wild-type nerves all showed near-linear amplitude decrement from minimal to maximal temperature, whereas *Fhf2*^*null*^ nerves displayed enhanced decrements above 41°C. (F) Histogram representation of CCAP amplitude decrements with increasing temperature. For each of 7 wild-type and 6 *Fhf2*^*null*^ nerve recordings, CCAP amplitude ratios for 32°C vs 28°C, 36°C vs 32°C, 40°C vs 36°C, and 44°C vs 40°C. The decrement in CCAP amplitude is significantly greater for *Fhf2*^*null*^ nerves vs wild-type nerves for 44°C vs 40°C. FHF, factor homologous factor.

The rapid reversibility of conduction deficit on nerve cooling is strong evidence that sodium channels in *Fhf2*^*null*^ axons at high temperature are not denatured or internalized but fail to generate sufficient inward current for impulse propagation due to accelerated inactivation. These findings parallel prior demonstration that hyperthermia-induced cardiac conduction block in *Fhf2*^*null*^ mice is ameliorated on cooling.^22^

### 3.5. Heat excitation deficit in a computational model of Fhf2^null^ nociceptor

To assess the requirement of FHF2 in mature heat-sensitive nociceptors, we used computational modeling and simulations. For these studies, the wild-type (*Fhf2*^*WT*^) nociceptor model was taken from the model cell described by Barkai et al.^4^ Their model has a highly branched terminal axon with uniformly distributed leak and voltage-gated channels except for the distal afferent terminals, which lack voltage-gated sodium channels but can generate inward currents mimicking the opening of TRP receptor channels.^4^ The compartmental segregation of excitatory currents from voltage-gated channels was based on recordings showing that action potentials initiate at recessed sites on nociceptive afferents.^16^ We simulated FHF2 loss-of-function by modifying the gating of the embedded TTX-sensitive and TTX-resistant conductances to reflect shifts in inactivation rates and voltage dependence commensurate with our recording data (**Fig. 2**) without altering conductance densities. All channel gating transitions in both the *Fhf2*^*WT*^ and *Fhf2*^*null*^ nociceptor models, including TTX-sensitive and TTX-resistant sodium conductances, were accelerated with increasing temperature by using Q10 rate scaling factors.

We first tested whether the *Fhf2*^*null*^ model axon suffers conduction block at elevated temperature, analogous to ex vivo conduction block (**Fig. 4**). Wild-type and *Fhf2*^*null*^ model axons were point stimulated by current pulse, and action potential conduction was monitored over a distance of 5 mm. The wildtype model axon conducted an action potential at either 37°C (**Fig. 5A**) or 43°C (**Fig. 5B**). By contrast, the *Fhf2*^*null*^ model axon conducted at 37°C (**Fig. 5C**) but suffered conduction block at 43°C (**Fig. 5D**).

**Figure 5.**
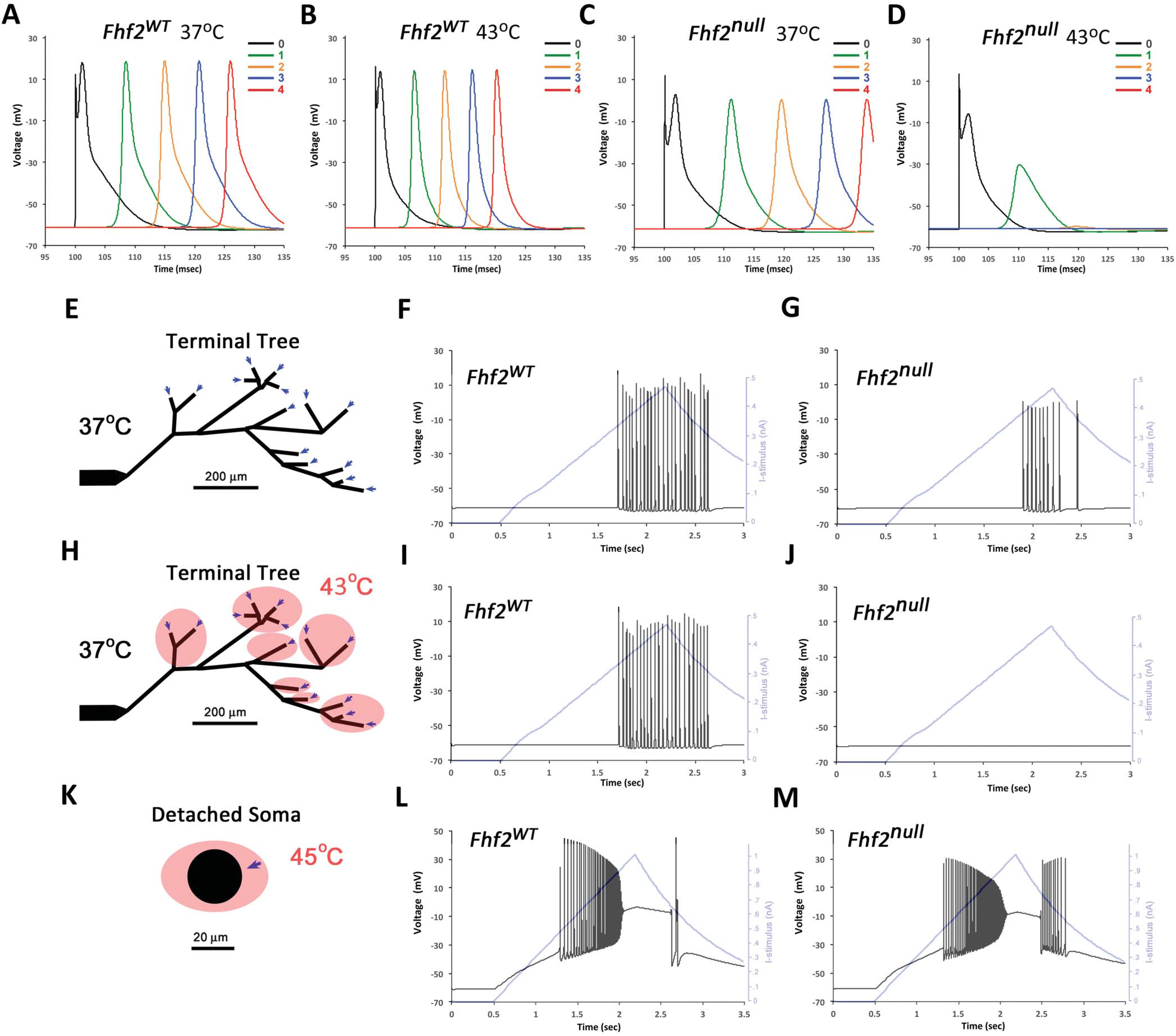
Computational modeling of wild-type and *Fhf2*^*null*^ nociceptors. Wild-type and *Fbf2*^*null*^ nociceptor models differ only in the inactivation gating parameters of the TTX-sensitive and TTX-resistant voltage-gated sodium conductances. All gated conductances have temperature-dependent state transition rates. (A-D) Simulation of axonal conduction. The long axon of each model was stimulated at one point with a 100 **m**s current pulse, and membrane voltage was monitored at 0, 1, 2, 3, and 4 mm from the site of stimulation. (A) Wild-type model at 37°C. (B) Wild-type model at 43°C. (C) *Fhf2*^*null*^ model at 37°C. (D) *Fbf2*^*null*^ model at 43°C; note conduction failure beyond 1 mm from the point of stimulation. (E-G) Simulation of afferent mechanical nociception. (E) Branching morphology of the afferent terminal. The terminal architecture, as published elsewhere,^4^ was incorporated into wild-type and *Fbf2*^*null*^ models. Temperature throughout nociceptorwasset to 37°C. Arrows at all terminal branch tips indicate sites of upramping current injection delivered from 0.5 to 2.2 seconds of simulation followed by downramping from 2.2 to 3 seconds. This protocol simulated opening of TRP channels at branch tips due to mechanical stimulation. (F) Central axonal voltage recording of wild-type model neuron during terminal stimulation. The current ramp (light blue trace) evoked a train of action potentials that initiated peripherally and propagated antidromically. (G) Voltage recording of *Fbf2*^*null*^ model neuron during terminal simulation. A train of action potentials was generated, although fewer in number than in the wild-type model. (H-J) Simulation of afferent heat excitability. (H) Terminal branching architecture. Branch segments within 200 μm of tips were set to 43°C (red), whereas remainder of the neuron was maintained at 37°C. Current ramp injection into branch tips simulated heat stimulation. (I) Central axonal voltage recording of wild-type neuron during terminal stimulation that evokes a train of antidromic action potentials. (J) Voltage recording of *Fhf2*^*null*^ neuron during terminal simulation. The mutant neuron was not excited by the stimulation of heated terminals, simulating the heat nociception deficit observed in vivo. (K-M) Simulation of heat-induced excitation in “acutely dissociated” model soma. (K) Model soma were disconnected from axons; somatic temperature was set to 45°C (red), and upramped somatic inward current simulated heat-induced excitation of the isolated soma. (L) Wild-type and (M) *Fhf2*^*nul1*^ isolated soma fired trains of action potentials in response to current injections (light blue traces). FHF, factor homologous factor; TRP, transient receptor potential; TTX, tetrodotoxin; WT, wild-type.

Simulations were next run by current injection into afferent terminals of nociceptor models at 37°C, which recapitulates the effects of noxious chemical or mechanical stimulation (**Fig. 5E**). The *Fhf2*^*WT*^ model fired a train of action potentials in response to such stimulation (**Fig. 5F**). The *Fhf2*^*null*^ nociceptor model also generated action potentials, albeit fewer in number, during current injection (**Fig. 5G**), consistent with preservation of mechanosensation (**Fig. 1C**) and chemosensation^35^ in *Fhf2*^*nul1*^ mice. Heat nociception was simulated by injecting current into afferent terminals, whereas the temperature of distal axon segments was elevated to 43°C while leaving the rest of the model neuron at 37°C (**Fig. 5H**). Under these conditions, the *Fhf2*^*WT*^ nociceptor model still generated a train of action potentials (**Fig. 5I**), whereas the *Fhf2*^*null*^ model failed to be excited (**Fig. 5J**).

In an attempt to explain why acutely dissociated *Fhf2*^*nul1*^ nociceptor neurons display relatively normal heat-induced excitation, we simulated acute dissociation of nociceptive neurons by severing the entire axons in the nociceptor models and performing simulated somatic current injection at 45°C (**Fig. 5K**). Under these conditions, both wild-type and *Fhf2*^*null*^ models generated a train of action potentials during current injection (**Figs. 5L and M**). These findings emphasize the importance of FHF2 to preserve sodium channel availability at an elevated temperature during passive propagation of depolarization from nociceptor axon terminal to trigger zone and during conduction down the axon.

## 4. Discussion

We have shown here that FHF2 is a powerful inactivation gating modulator for the principle TTX-sensitive and TTX-resistant sodium channel isoforms expressed in nociceptive neurons. FHF2 induces depolarizing shifts in V_1/2_ inactivation for Na_v_1.7 and TTX-resistant sodium channels and slows the rate of Na_v_1.7 inactivation from closed and open states. Elevating temperature acts as an independent accelerant of inactivation, speeding closed and open state Na_v_1.7 inactivation regardless of the FHF2 association. Hence, Na_v_ sodium flux on membrane depolarization is most compromised at elevated temperature in the absence of FHF2. These gating principles offer a plausible explanation for the specificity of heat nociception deficit in *Fhf2*^*null*^ mice, despite the broad expression of FHF2 in all classes of sensory neurons. These principles also provide a rationale for the preservation of mechanosensation in *Fhf2*^*null*^ mice, which is mediated, at least partially, by the same nociceptors that respond to noxious temperatures.^30^

Loss of heat nociception was observed following either global or peripheral neuron-restricted genetic ablation of the *Fhf2* gene, demonstrating the requirement of FHF2 function in heat-sensitive nociceptors. We were initially surprised to find that heat-induced excitability of acutely dissociated *Fhf2*^*null*^ DRG neurons was unimpaired. This paradox is likely explained by the fact that acutely dissociated neurons lack axonal processes and are electrotonically compact; with heat-inducible TRP channels and voltage-gated channels comingled on the somatic plasma membrane, inward TRP channel currents directly depolarize the somatic membrane bearing voltage-gated sodium channels. By contrast, depolarization of heated nociceptor afferent terminals produces retrograde axial current that must passively depolarize a periterminal trigger zone^16^ subject to cable property delay, reducing sodium channel availability in the absence of FHF. Furthermore, reduced sodium current duration in the *Fhf2*^*null*^ afferent may provide insufficient charge to ensure spike conduction, consistent with our observed failure of unmyelinated *Fhf2*^*null*^ sensory axons to sustain C-fiber compound action potentials on heating. Our computational modeling further supports this hypothesis. Although current influx at afferent terminals of the *Fhf2*^*null*^ model neuron failed to evoke action potentials at 43°C, simulated injection of current into the heated axon-severed soma was able to trigger excitation. It is also possible that axotomy accompanying preparation of acutely dissociated DRG neurons triggers upregulation of channel expression to provide sufficient sodium current in mutant cells at elevated temperature.

Although peripheral sensory axons express FHF2, myelinated and unmyelinated axons in the central nervous system show little or no FHF expression.^10^ This natural FHF deficiency may explain why spike conduction in brain axons is blocked under hyperthermal conditions.^24^

We used plasmid-transfected HEK293 cells to assay the effects of FHF2B expression on the inactivation of Na_v_1.7. The voltage dependence of Na_v_1.7 steady-state inactivation was markedly right shifted by 15 mV from V_1/2_ of −87 mV in the absence of FHF2B to −72 mV in its presence. As discussed above, strong modulation of Na_v_ inactivation by FHF2 is central to the heat-associated nociception and cardiac deficits in *Fhf2*^*nul1*^ mice. Our findings are in sharp contrast to Yang et al.^35^ who reported only a minor shift in Na_v_1.7 V_1/2_ inactivation from −76 mV in HEK cells without FHF cotransfection to −73 mV in presence of FHF2B. These disparate findings may reflect expression of endogenous FHF2 protein in some sublineages of HEK293 cells that result in a significant depolarizing shift in Na_v_1.7 V_1/2_ inactivation in the absence of FHF transfection. Alternatively, the expression vector used by Yang et al. may have produced a lower concentration of FHF2 insufficient to saturate binding to Na_v_1.7. It is possible that the failure by Yang et al. to detect a strong inactivation modulatory effect of FHF2 led them to emphasize other mechanistic bases for the *Fhf2*^*nul1*^ heat nociception deficit.^35^ Yang et al.’s proposal that FHF2 prevents partial internalization of Na_v_1.7 at elevated temperature is not substantiated by our finding that Na_v_1.7 peak current is relatively stable at 40°C in the absence of FHF (**Fig. 2I**).

FHF2 deficiency causes hyperthermia-induced cardiac conduction block^22^ along with the heat nociception deficit reported here and elsewhere.^35^ Our findings show that these disparate phenotypes share a common underlying molecular mechanism. FHF2 increases the availability of cardiac and peripheral neuronal sodium channels at rest by raising the voltage of steady-state channel inactivation. Furthermore, FHF2 delays the inactivation of these channels from closed and open states, thereby counteracting heat-induced acceleration of inactivation and thus allowing for sufficient sodium current to drive action potential conduction through heated nociceptive axons or hyperthermic myocardium.

## 5. Conclusions

FHF2 expression in sensory neurons is absolutely essential for heat nociception. FHF2 modulation of TTX-sensitive and TTX-insensitive sodium channel inactivation gating allows for sufficient inward sodium currents on heat-induced depolarization to enable axonal initiation and propagation of antidromic action potentials. This conclusion is supported by voltage-clamp analysis of sodium currents, by C-fiber action potential conduction, and by initiation and conduction of action potentials in a nociceptive neuron computational model as a function of temperature and FHF2 expression status. The heat nociception deficit in *Fhf2*^*null*^ mice is mechanistically equivalent to cardiac conduction deficit when these mice are exposed to core hyperthermia.

## Acknowledgments

The authors thank Sergio Solinas (University of Sassari, Sardinia) for expert assistance in implementing modifications to the nociceptor neuron computational model of Barkai et al.^4^ The authors thank Glenn I. Fishman (NYU Langone Medical Center) for his encouragement and support. This work was supported by R01HL142498 to G.I. Fishman and M. Goldfarb, a CUNY research grant to M. Goldfarb, and the Blaustein Pain Research Foundation and the Neurosurgery Pain Research Institute at Johns Hopkins (to M. Ringkamp).

